# An overview on the DNA nucleotide compositions across kingdoms

**DOI:** 10.1101/087569

**Authors:** Yabin Guo

## Abstract

The DNA nucleotide compositions vary among species. This fascinating phenomenon has been studied for decades with some interesting questions remaining unclear. Recent years, thousands of genomes have been sequenced, but general evaluations on the nucleotide compositions across different phylogenetic groups are still absent. In this letter, I analyzed 371 genomes from different kingdoms and provided an overview on DNA nucleotide compositions. A number of important topics were discussed, including GC content, DNA strand symmetricity, CDS purine content, codon usage, thermophilicity in prokaryotes and non-coding RNA genes. I also gave explanations to two long debated questions: 1) both genome GC content and CDS purine content are correlated with the thermophilicity in archaea, but not in bacteria; 2) the purine rich pattern of CDS in most species is mainly a consequence of coding requirement, but not mRNA interaction dynamics. This study provides valuable information and ideas for future investigations.

## Main text

The DNA molecules in all organisms are composed of the same four nucleotides, A, T, G and C, while the ratios of the four nucleotides vary among species, which has been fascinating to people for nearly a century. In 1950s, Erwin Chargaff found that in DNA the number of G equals the number of C, and the number of A equals the number of T, which is known as the Chargaff’s first rule (Chargaff et al. 1950; Chargaff et al. 1952). Now we know that it is correct in double strand DNA for the complement between purines and pyrimidines. In 1960s, Chargaff published his second rule (the second parity rule, PR2), which stated that in each DNA strand the A ratio roughly equals the T ratio, and the G ratio roughly equals the C ratio (Rudner et al. 1968). This rule has been proved largely true except in some small DNA molecules such as the mitochondrial (mt) DNAs. Beside the overall genomic nucleotide composition, the nucleotide composition of coding sequences (CDS) is also an important topic. Szybalski et al. (Szybalski et al. 1966) and Smithies et al. (Smithies et al. 1981) found that DNA template strands have more pyrimidine nucleotides (i.e. RNAs are purine rich), which was later named the Szybalski’s rule by Forsdyke (Dang et al. 1998; Lao and Forsdyke 2000). Forsdyke claimed that Thermophiles strictly obey Szybalski’r rule and raised a *Politeness Hypothesis*, assuming that mRNA with higher purine content are “polite” to avoid undesired interactions, and mRNA of thermophiles need to be even more polite, because the entropy-driven reactions are more prone to happen under high temperature (Lao and Forsdyke 2000). However, the results of further studies turned out to be paradoxical (Paz et al. 2004; Mahale et al. 2012). So far, the applicability of Szybalski’s rule has not been proved.

Most of these studies were performed in the *pre-genomics era* and sometimes based on incomplete genomic data. During the recent ten years, thanks to the development of next generation sequencing technology, genomes of thousands of species were sequenced. Yet, there still lacks a global evaluation on the DNA nucleotide compositions across kingdoms (or domains). In this letter, I analyzed 371 genomes (122 animals, 39 plants, 53 fungi, 32 protists, 25 archaea and 100 bacteria) and revealed a number of amazing facts and provided explanations for two long unsolved questions.

First, the GC contents of all the nuclear genomes were calculated (Fig. 1A, Table S1). The GC contents of animal genomes have the smallest diversity with an average of 40%, and more invertebrate genomes have lower GC contents than vertebrate genomes do. Most of the plant genomes analyzed here falls into two groups: the dicots (yellow fill) with lower GC contents and the grass family (Poaceae) monocots (blue fill) with higher GC contents (Kumari and Ware 2013; Smarda et al. 2014). Banana (*M. acuminata*), the only non-Poaceae monocot analyzed (red fill) has a medium GC content between the two groups. There are three plant genomes have considerably higher GC contents. Actually, they are green and red algae instead of Embryophytes. Protists and prokaryotes are more complex phylogenetic groups and it is not surprising that their genome GC contents have higher diversity. Among all known genomic sequences, bacterium, *Anaeromyxobacter dehalogenans*, has the highest GC content (74.9%), while *Candidatus* Zinderia insecticola (a symbiont in spittlebugs) has the lowest GC content (13.5%), even lower than all known mitochondrial genomes (Nishida 2013). Archaea have relatively moderate DNA GC contents compared with bacteria, though many of them live in extreme environments. The genome of *Plasmodium falciparum* (one of the malaria parasite) has the lowest GC content (19%) in all eukaryotic genomes (Gardner et al. 2002).

**Fig. 1.**
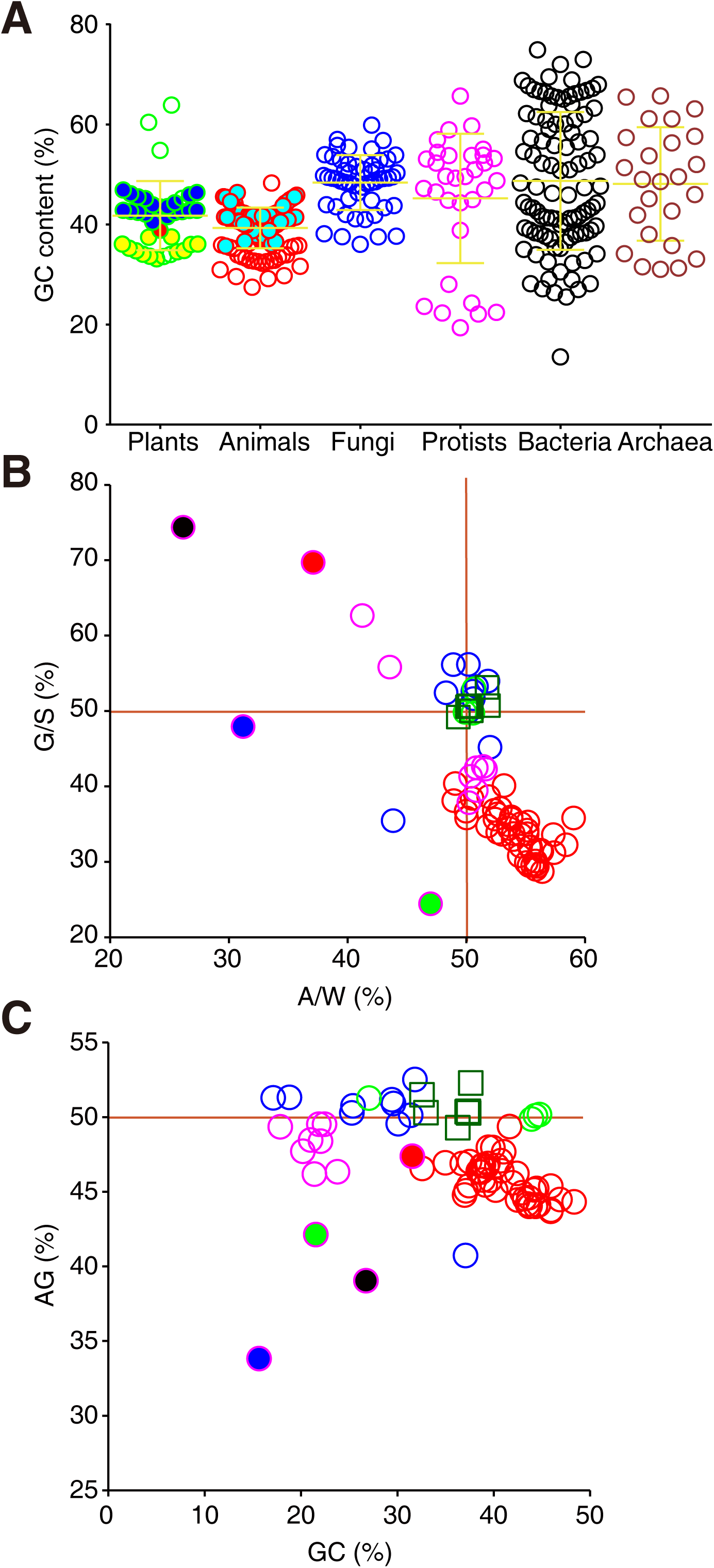
Genome GC contents and mitochondrial nucleotide compositions. Each point is one species. A, genome GC contents across kingdoms (blue fill, plants of Poaceae; yellow fill, dicot plants; red fill, *Musa acuminata*; cyan fill, vertebrates); B, C, G/S-A/W (B) and AG-GC (C) plots for genomes of mitochondria and chloroplasts (red: vertebrates; magenta, invertebrates; green, plants; blue, fungi; dark green square, chloroplasts/plastids; blue fill, *Mnemiopsis leidyi*; green fill, *Atta cephalotes*; red fill, *Schistosoma mansoni*; black fill, *Onchocerca volvulus*).

Then, the Chargaff’s second parity rule was evaluated. All large chromosomes are symmetric as expected (Table S1). Whereas, many mtDNAs have asymmetric strands as described previously (Francino and Ochman 1997; Frank and Lobry 1999). All animal mitochondrial genomes are small (10-20 kb, Fig. 1B, Table S1). The invertebrate mtDNAs have lower GC contents (21-32%) than those of vertebrates (32-49%). Although tunicate (*Ciona intestinalis*) is chordate and evolutionally much closer to vertebrates than to protostomes, its mtDNA nucleotide composition is more similar to those of protostomes (Fig. 1C). In a typical vertebrate mtDNA, one strand has more A and C, while the other stand has more G and T. The nucleotide compositions of invertebrate mtDNA have large diversity. The mtDNAs of leaf-cutting ant (*Atta cephalotes*, green fill), comb jelly (*Mnemiopsis leidyi*, blue fill), blood fluke (*Schistosoma mansoni*, red fill) and river blindness parasite (*Onchocerca volvulus*, black fill) are scattered far away from the cluster, matching their special places in evolution (Fig. 1B, C). The plant mtDNAs, chloroplast DNAs and fungal mtDNAs are usually larger and have more symmetric strands, distributing around point (50, 50) in the G/S-A/W plot (Fig. 1B).

To indicate how far a DNA strand is from symmetry more delicately, an index *Asy* (asymmetricity) is introduced (see Methods section). Briefly, *Asy* is the Euclidean distance between the point of a given DNA strand and the point (50, 50) on the G/S-A/W plot. Consisting with previously reported, the small mtDNAs have higher *Asy* values, but there is no correlation between *Asy* and mtDNA size (Fig. S1A). Among all the mtDNA analyzed, *O. volvulus* has the most asymmetric strands. Notably, the mtDNA of *Puccinia graminis* (the fungal pathogen of black rust in wheat) has a pretty high *Asy*, though its size is many times larger than typical animal mtDNAs (magenta fill in Fig. S1A, Table S1).

Small perturbations are found in the *Asy* values of prokaryotic and protist chromosomes, which is not surprising for their small sizes. And protist chromosomes show larger diversity in *Asy* than prokaryotic chromosomes do when their sizes are similar. One third of the Leishmania chromosomes have considerably high *Asy* values (Fig. S1B). Moreover, the G ratio correlates well with the A ratio in Leishmania chromosomal sequences, indicating there are heavy and light strands (Fig. S1C).

Statistically, larger chromosomes tend to have more symmetric strands than smaller ones do, while larger chromosomes also tend to have more asymmetric local regions. Indeed, the unassembled scaffolds/contigs of animal genomes show a substantially scattered pattern on the G/S-A/W plot (Fig. S2A). For example, in a 1.86 Mb scaffold of the kangaroo rat (*Dipodomys. ordii*) genome, the number of A is 8 times of that of T (*Asy*=40.5). These unassembled scaffolds/contigs usually are highly repeated and many of them contain satellite DNA located in centromeres and telomeres. It is known that satellite DNA comprises more than half of the *D. ordii* genome (Mazrimas and Hatch 1972). Besides *D. ordii*, large contigs with asymmetric strands can also be found in dog (*C. familiaris*) and collared flycatcher (*Ficedula albicollis*) genomes (Fig. S2B). Although many plant genomes contain large fractions of repetitive regions, similar highly asymmetric regions were not found in plant genomes (Fig. S2B).

Symmetry (high entropy) is more stable than asymmetry (low entropy). Inversions, translocations and transpositions make DNA strands more and more symmetric as time goes by (Albrecht-Buehler 2006). Asymmetric strands in chromosomal regions (or small chromosomes) may be maintained by special replication mechanisms (e.g. for mtDNAs) or evolutionary benefits (e.g. for satellite DNAs), but it is impossible for a large chromosome to maintain asymmetric strands due to the high energy barrier. Actually, it is not surprising that Chargaff’s second rule is correct, and it would be really surprising if it is not correct.

To test Szybalski’s rule, I calculated the CDS of all the genomes mentioned above (Table S2). The GC contents of CDS correlate well with the GC contents of genomes, especially in prokaryotes, because CDS comprise most of the prokaryotic genomes, while in eukaryotes, the GC contents of CDS usually are greater than those of genomes (Fig. S3A). G/S-A/W plot shows that animal, plant and fungal CDS distribute in three different areas, while protist and prokaryotic CDS show large diversities (Fig. 2A). Similar to the previous reports in prokaryotes (Lao and Forsdyke 2000; Mahale et al. 2012), AG contents are negatively correlated with GC contents among all species (R^2^=0.497, Fig. 2B). The average purine contents of CDS (APCC) of plant, animal and protist genomes are all greater than 50%. However, fungal genomes have relatively higher GC contents (Fig. 1A) and lower APCC. 15 of the 53 fungal APCC are less than 50%. All the 25 archaeal APCC are greater than 50%, whereas, 22 of 100 bacterial APCC are less than 50%. Therefore, Szybalski’s rule is not well complied in certain organisms (Fig. 2B, Fig. S3B, Table S2).

**Fig. 2.**
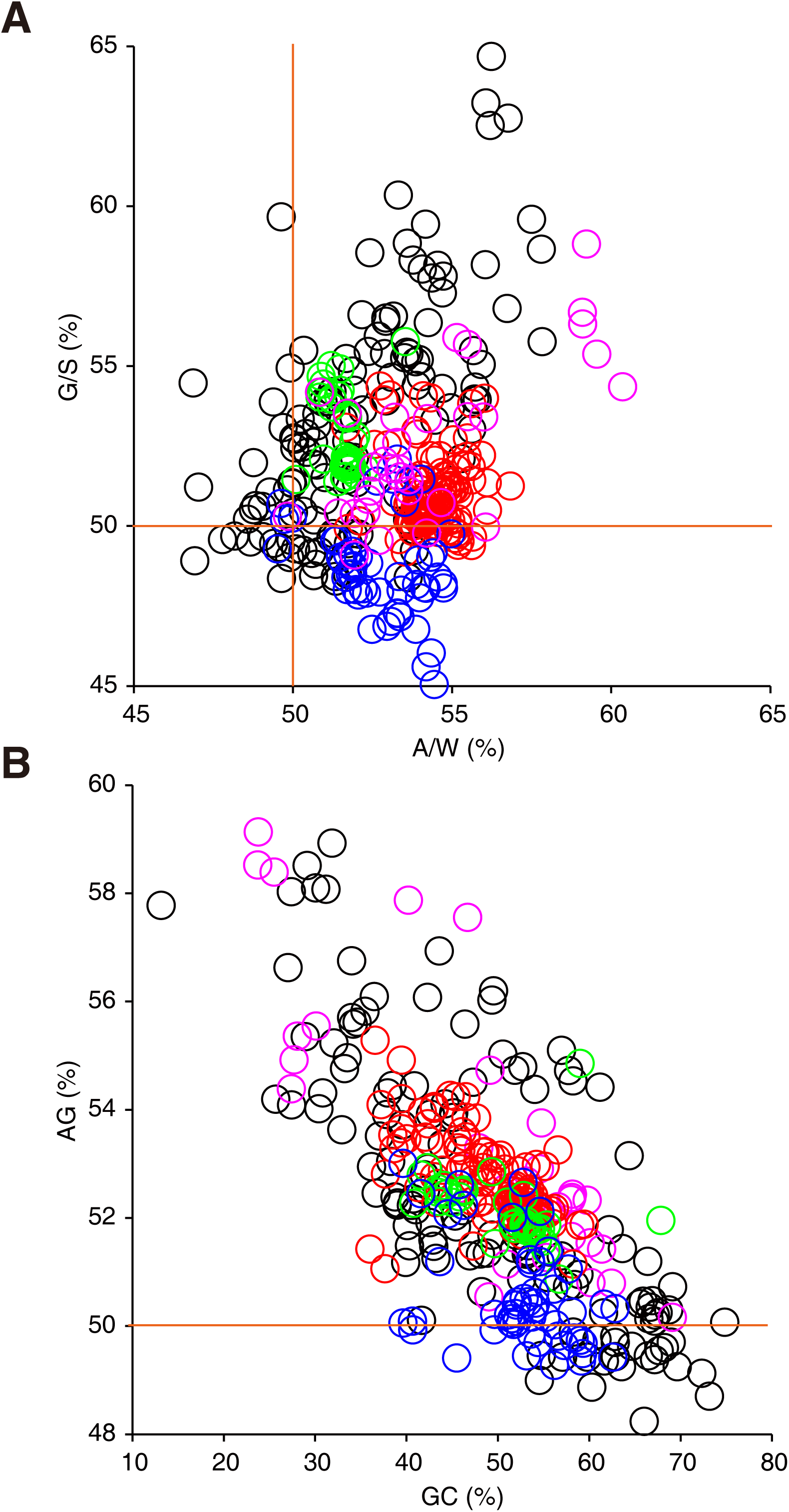
Nucleotide compositions of CDS. A, G/S-A/W plot; B, AG-GC plot (red: animals; green, plants; blue, fungi; magenta, protists; black, prokaryotes).

The influence of genome GC content (Kagawa et al. 1984; Galtier and Lobry 1997; Musto et al. 2004; Wang et al. 2006) or CDS purine content (Lao and Forsdyke 2000; Paz et al. 2004; Mahale et al. 2012) on the thermophilicity in prokaryotes has long been debated and remained unclear. Fig. S4 showed that APCC negatively correlate with the genome GC contents in both archaea and bacteria as previously described (Lao and Forsdyke 2000; Mahale et al. 2012). Notably, 12 of 14 thermophilic archaea are above the trend line and all mesophilic archaea are under the line, which strongly suggests that both purine and GC contents contribute to thermophilicity (Fig. S4A). However, similar phenomenon was not observed in bacteria. Many mesophilic bacteria are above the trend line, for example, the three species with APCC>58% are all pathogens (Fig. S4B). The controversial observation in previous studies could be partially due to that archaea and bacteria were not treated separately. More importantly, since both purine and GC contents contribute to thermophilicity, it is hard to predict thermophilicity with just one value of them. According to the equation of the trend line (Fig. S4A), a new index *TI* (Thermo Index) was introduced: *TI=AG%+0.14GC%*. To test the applicability of *TI*, another batch of archaeal genomes were analyzed (Table S3). Thermophiles and mesophiles can be roughly distinguished by using CDS purine contents (Fig. S5A, B), but not genome GC contents (Fig. S5C, D), whereas they are distinguished far more delicately by using *TI* (Fig. S5E, F). These results indicate that there is a tradeoff between purine and GC contents, and the two values should be considered together with well-balanced weights to provide accurate predictions for thermophilicity (see detailed discussion in supplementary text).

Previous studies also showed that halophile genomes have high GC content (Paul et al. 2008). In this study, all the halophilic archaea have GC content>60%, and on the other hand, all the archaea with GC content>60% are halophiles, including *Methanopyrus kandleri*, which is both thermophilic and halophilic (Fig S4A). Halophilic bacterium, *Salinibacter ruber*, has 66% GC in its genome as expected, while a number of bacteria with GC content>60% are not halophilic (Fig. S4B).

The amino acid (AA) frequencies and the codon usages were also calculated (Fig S6). Prokaryotes show a distinct AA frequency profile from eukaryotes. The most striking fact is that prokaryotes have higher frequencies than eukaryotes do in all aliphatic AAs (Gly, Ala, Val, leu and Ile), as well as much lower frequencies in serine and cysteine. Cysteine frequencies are notably higher in animals than in other groups (Fig. S6A, B). If looking at the individual genomes, the most extreme ones are malaria parasites. With extremely low GC contents, some members in *Plasmodium* genus have very high frequencies of Asn (Fig. S6B), Tyr and Lys and very low frequecies of Leu, Pro, Arg, Val, Ala and Gly. AAT comprises more than 12% codons in *P. falciparum* CDS (Fig. S6C).

The nucleotide compositions of the three positions of codons were characterized respectively (Fig. 3). Position 2 shows the smallest diversity, especially T2 and G2, while position 3 shows the largest diversity. Some species can even have only 1-3% A, T, G or C in position 3 (Fig. 3A). In the G/S-A/W plot, the three positions fall into three distinct groups, no matter what kingdom one species belonging to, which strongly suggests that the nucleotide compositions of CDS are largely constrained by the coding requirement (Fig. 3B). The purine content of position 1 in all species is significantly greater than 50%, while position 2 and 3 basically have more pyrimidine nucleotides. Obviously, high APCC is mainly contributed by position 1. Since position 3 is not constrained by the amino acid frequencies, it is the best reflection of selection pressure in mRNA interaction dynamics. If the *Politeness Hypothesis* were true, position 3 should have been the major contribution of the purine rich pattern of mRNAs, but actually its average purine content is less than 50% in most species. Therefore, it seems that mRNAs tend to be slightly pyrimidine rich, whereas, position 1 can only be virtually purine rich, because all three stop codons and codons for some low-content amino acids (Cys, Trp, Tyr, His) are started with pyrimidines.

**Fig. 3.**
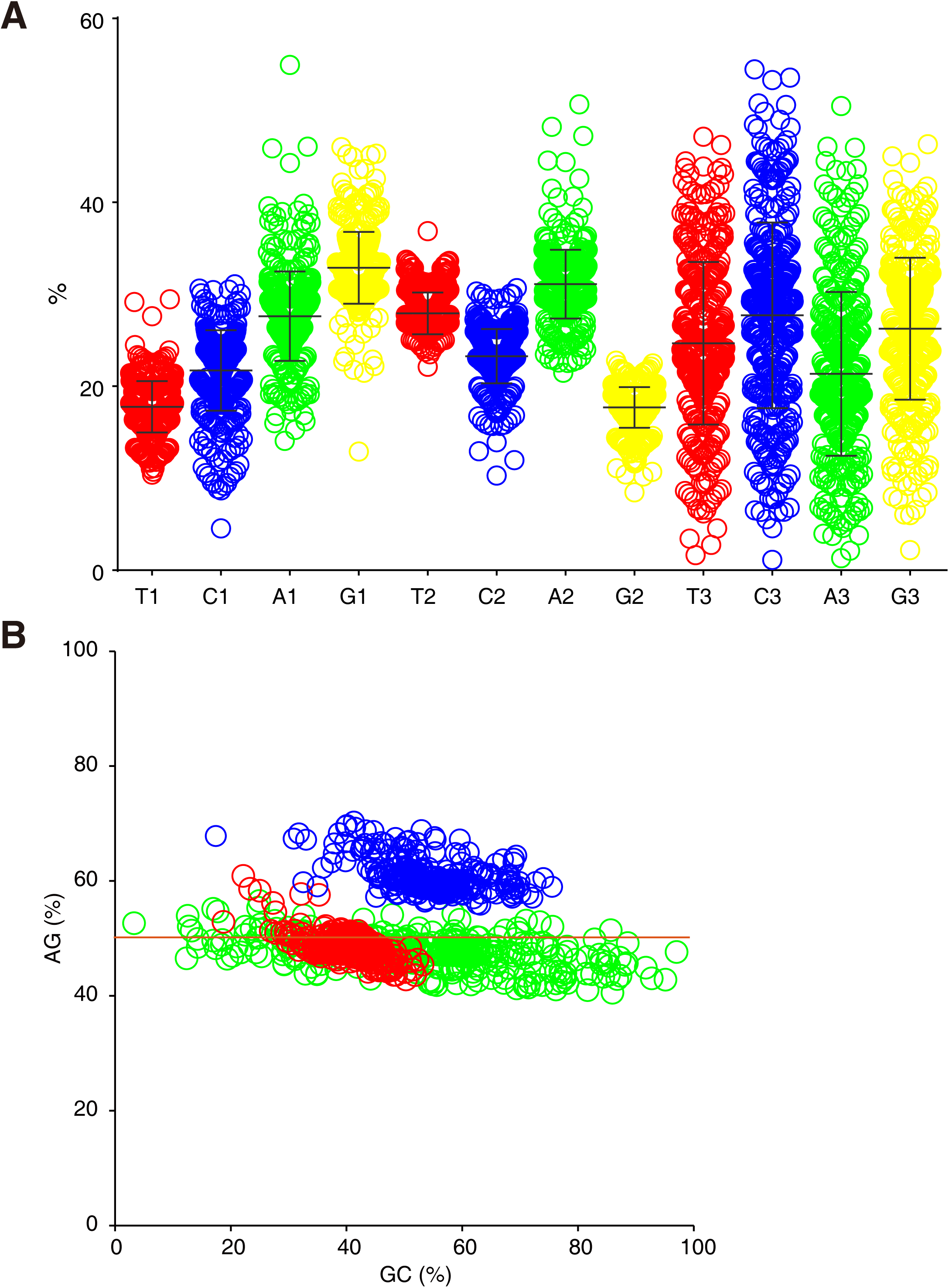
Nucleotide compositions of the three positions of codons. A, percentages of T, C, A and G of the three positions (red, T; blue, C; green, A; yellow, G); B, AG-GC plot of the three positions of codons (blue: position 1; red: position 2; green: position 3).

Finally, I calculated the nucleotide compositions of untranslated regions (UTR), introns and noncoding RNA (ncRNA) genes (if available) (Table S4). Similar to CDS, the GC contents of UTR and introns are well correlated with the GC contents of genomes, but the correlation between the GC contents of ncRNA genes and the GC contents of genomes is weak (Fig. S7A, B). Unlike CDS, these sequences are not purine rich, though they are all transcribed units. Also, there’s no correlation between the AG contents and GC contents in these regions (Fig. S7C, D). The purine contents of ncRNA genes in thermophilic archaea are not higher than those in mesophilic archaea. Interestingly, the GC contents of ncRNA genes in thermophiles are significantly higher than those in mesophiles (Fig. S8). The function of mRNAs is mainly based on primary structure; whereas the function of ncRNAs is heavily rely on secondary structure. Therefore, it is not surprising ncRNAs have distinct nucleotide composition pattern from mRNAs even under the same environment. These results also suggest that the purine rich pattern and tradeoff between purine and GC contents in CDS are mainly a consequence of coding requirement.

Considering that purine synthesis consumes more energy than pyrimidine synthesis, CT rich RNAs could be an advantage in evolution for saving energy, which may partially explain why introns tend to be CT rich. Therefore, there should be a balance between genome GC content, coding requirement, energy optimization and interaction dynamics, which shapes the nucleotide composition of CDS. The nucleotide composition is evolutionarily flexible. Environment can have great influence (Foerstner et al. 2005), but obviously it can’t account for all the results. The actual situation is far more complicated than what we currently know. The determinants of nucleotide composition are still to be answered by future researches.

By analyzing 371 genomes (including genomic sequences, CDS, UTR, introns and ncRNAs), this letter provided an overview on the DNA nucleotide compositions across kingdoms, and showed that: 1) Chargaff’s second rule is largely true. 2) Some protists, such as Leishmania have relatively asymmetric chromosomal strands. 3) Long asymmetric satellite DNA regions were found in some animal genomes, but not in plant genomes. 4) Szybalski’s rule is not universal. CDS in certain bacterial and fungal genomes tend to be pyrimidine rich. Non-coding transcribed regions don’t comply with Szybalski’s rule in any phylogenetic groups. The purine rich pattern in CDS is mainly a consequence of coding requirement. 5) Thermophilicity and halophilicity are well correlated with nucleotide compositions in archaea, but not in bacteria. The previously controversial observations were mainly because that the genome GC contents and CDS purine contents were not considered together.

Additionally, this study also provided valuable datasets that researchers can use conveniently in future studies.

## Methods

(see Supplementary Materials)

## Supplementary materials

### Methods

#### Data resource

All sequence files (FASTA) were obtained from the Ensembl ftp sites (ftp.ensembl.org/pub/release-83/fasta for vertebrates; ftp.ensemblgenomes.org/pub/release-30/metazoa/fasta for invertebrates; ftp.ensemblgenomes.org/pub/release-30/fungi/fasta for fungi; ftp.ensemblgenomes.org/pub/release-30/plants/fasta for plants; ftp.ensemblgenomes.org/pub/release-30/bacteria/fasta for prokaryotes). Genome sequences are from .dna_sm_toplevel files; CDS are from .cds.all (eukaryotes) or .cdna.all (prokaryotes) files; noncoding RNAs are from .ncrna files; UTRs and introns are from .cdna.all files.

#### Equation for Asymmetricity (*Asy*)

The calculation of *Asy* based on G/S-A/W plot.

(*X, Y*) is a point on *G/S-A/W* plot.

*S=G+C; W=A+T*.

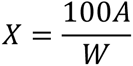

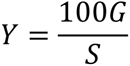

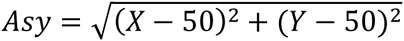

#### Programming

All scripts were written in Perl language.

### Supplementary text

#### G/S-A/W plot

Some studies use GC-skew and AT-skew to indicate the symmetricity of G~C and A~C. GC-skew=(G-C)/(G+C); AT-skew=(A-T)/(A+T). According to mathematical habit, formulas should be as simple as possible. Therefore, in this study, G/(G+C) and A/(A+T) are used instead of skews. The interval for G/S or A/W is [0,100%] (S=G+C; W=A+T), while the interval for skews is [-100%, 100%]. In one word, G/S-A/W is equivalent to skews, but with simpler form.

#### *TI* (Thermo Index) and thermophilicity in archaea

Because the interaction between G and C is stronger than the interaction between A and T, some researchers hypothesized that thermophiles would have higher GC content to make the secondary structure of their nucleic acids more stable. However, positive correlations were not found between GC content and thermophilicity in prokaryotes, and even negative correlation were found in some cases, so that it seems that thermophiles tend to have low GC contents. The reason for this confusing situation is mainly because thermophilicity is positively correlated with the CDS purine content in archaea, but CDS purine content is negatively correlated with genome GC content. Also, the contribution of CDS purine content weighs far more than the contribution of genome GC content. Therefore, if GC content is considered independently (without purine content being considered), the illusion that GC content negatively contributes to thermophilicity will occur. In Fig. S4A, most thermophilic archaea are above the trend line, indicating that the contributions of both purine content and GC content are positive. That is to say, if a thermophile and a mesophile have the similar GC contents, then the former is likely to have higher CDS purine content than the latter does; or if a thermophile and a mesophile have similar CDS purine contents, then the former is likely to have high GC content than the latter does. There is a tradeoff between CDS purine content and genome GC content, one contribution being compensated by the other. It would be quite unusual for a mesophile to have both higher GC content and higher CDS purine content than a thermophile does. Therefore a new index *TI* (Thermo Index) is introduced by balancing the contributions of CDS purine content and genome GC content based on the equation of the trend line in AG-GC plot. In the *TI* formula (*TI=AG%+0.14GC%*), both of the two values contribute to thermophilicity with different weights. *TI* has been proved so far to be applicable in distinguishing thermophiles from mesophiles in archaea (Fig. S5). Notably, there is no good correlation between thermophilicity and purine or GC content in bacteria. Considering the relation between bacteria and archaea is even farther than the relation between archaea and eukaryotes, it is not surprising that bacteria may have very different strategies from archaea in dealing with high temperature.

### Supplementary Tables

**Table S1.** Nucleotide compositions of nuclear genomes and mitochondrial/chloroplast/plastid genomes (if available).

**Table S2.** Nucleotide compositions, codon usages and amino acid frequencies of CDS.

**Table S3.** Nucleotide compositions of 10 mesophilic and 10 thermophilic archaea not including in Table S1 or S2 for testing the applicability of *TI*.

**Table S4.** Nucleotide compositions of UTR, introns and noncoding RNAs.

### Supplementary figure legends

**Fig. S1.**
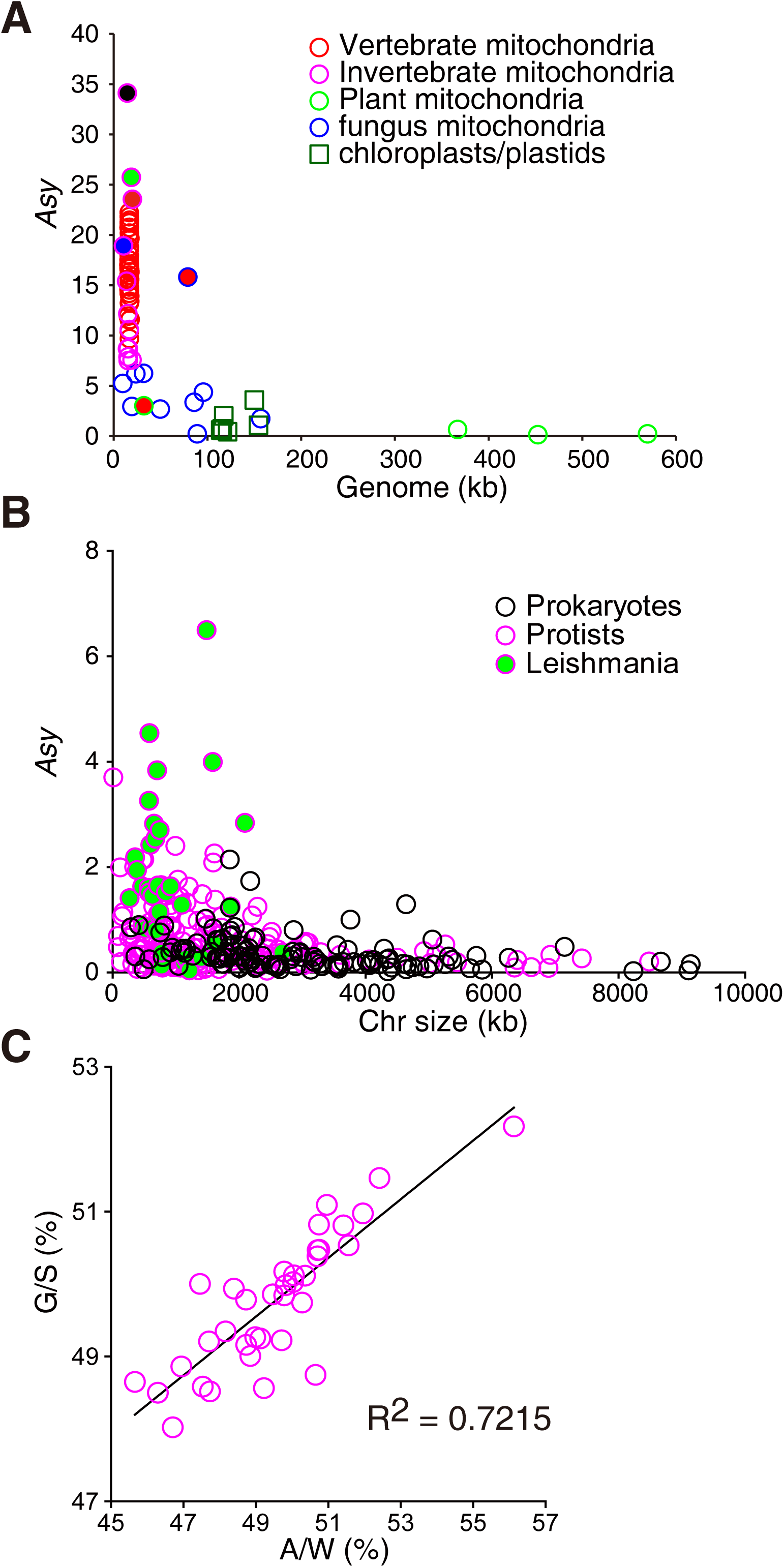
Strand asymmetricity of mtDNAs, prokaryotic chromosomes and protist chromosomes. A, strand asymmetricity of chromosomes of mitochondria and chloroplasts/plastids; B, strand asymmetricity of chromosomes of protists and prokaryotes, each point representing one chromosome; C, G/S-A/W plot of Leishmanial chromosomes.

**Fig. S2.**
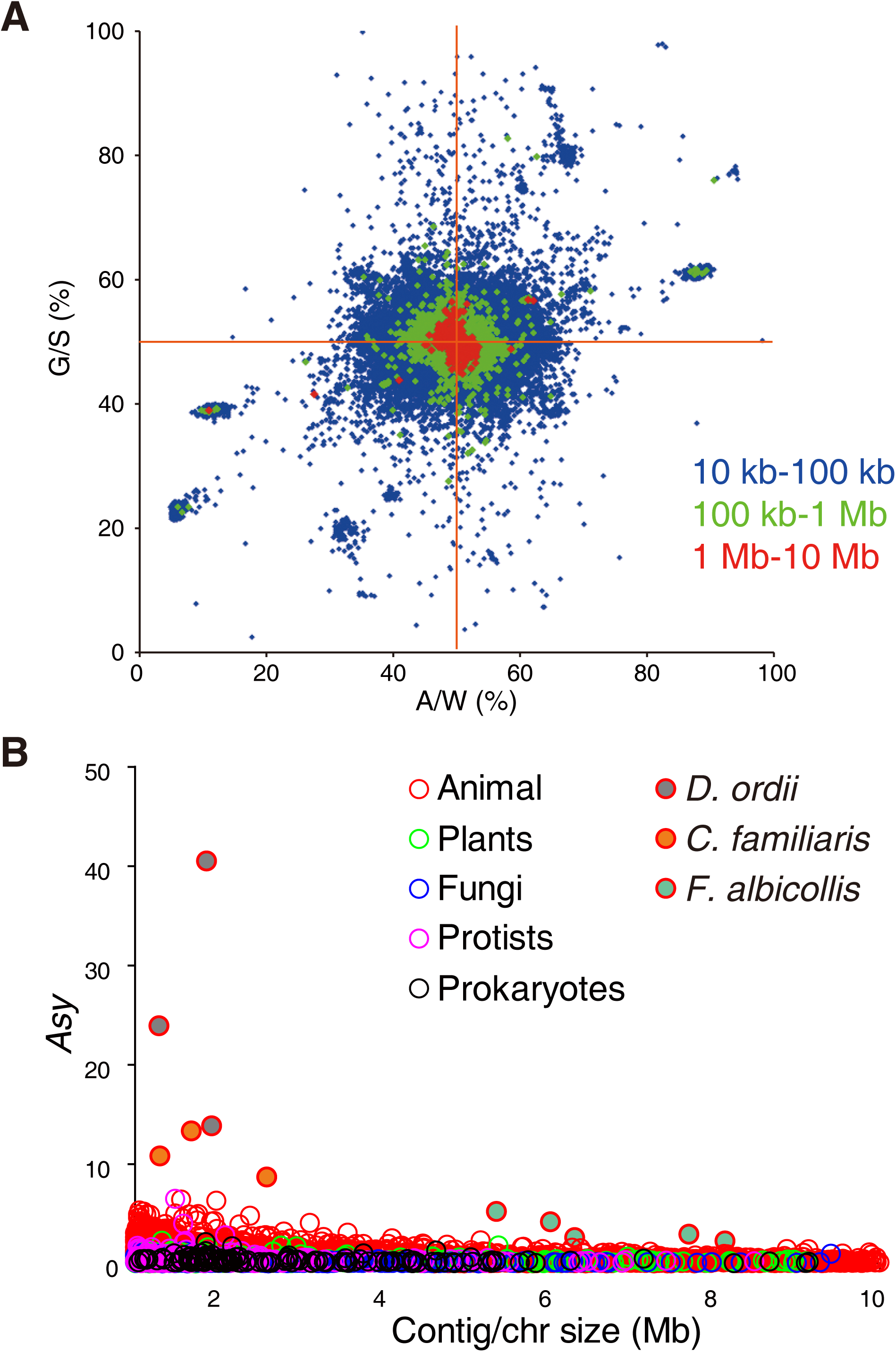
Strand asymmetricity of small chromosomes and contigs. A, G/S-A/W plot of contigs of animal genomes; B, strand asymmetricity of chromosomes/contigs less than 10 Mb.

**Fig. S3.**
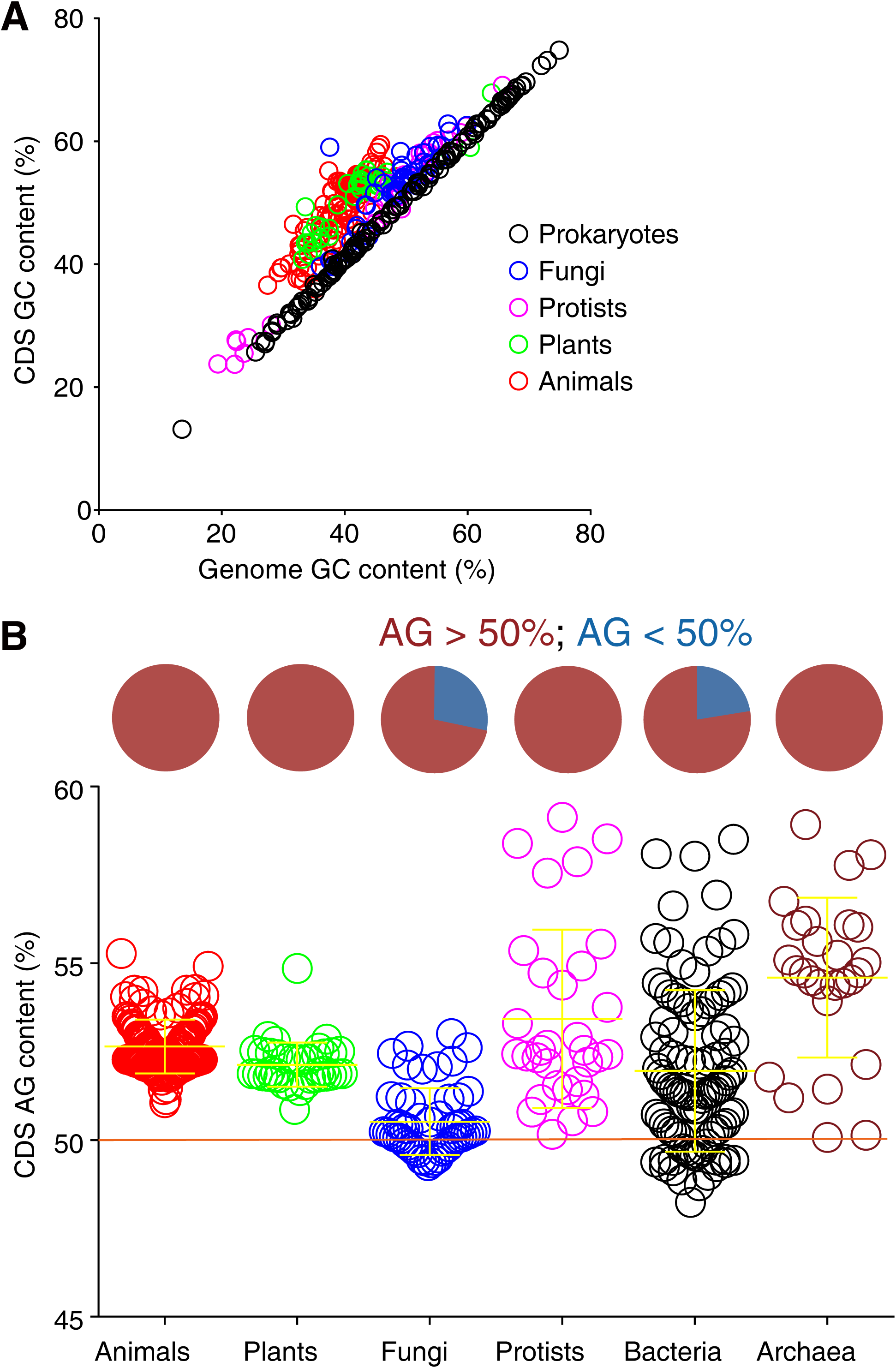
Nucleotide compositions of CDS. Each point is one species. A, the correlation between the GC contents of CDS and the GC contents of genomes; B, purine contents of CDS (APCC) of different groups.

**Fig. S4.**
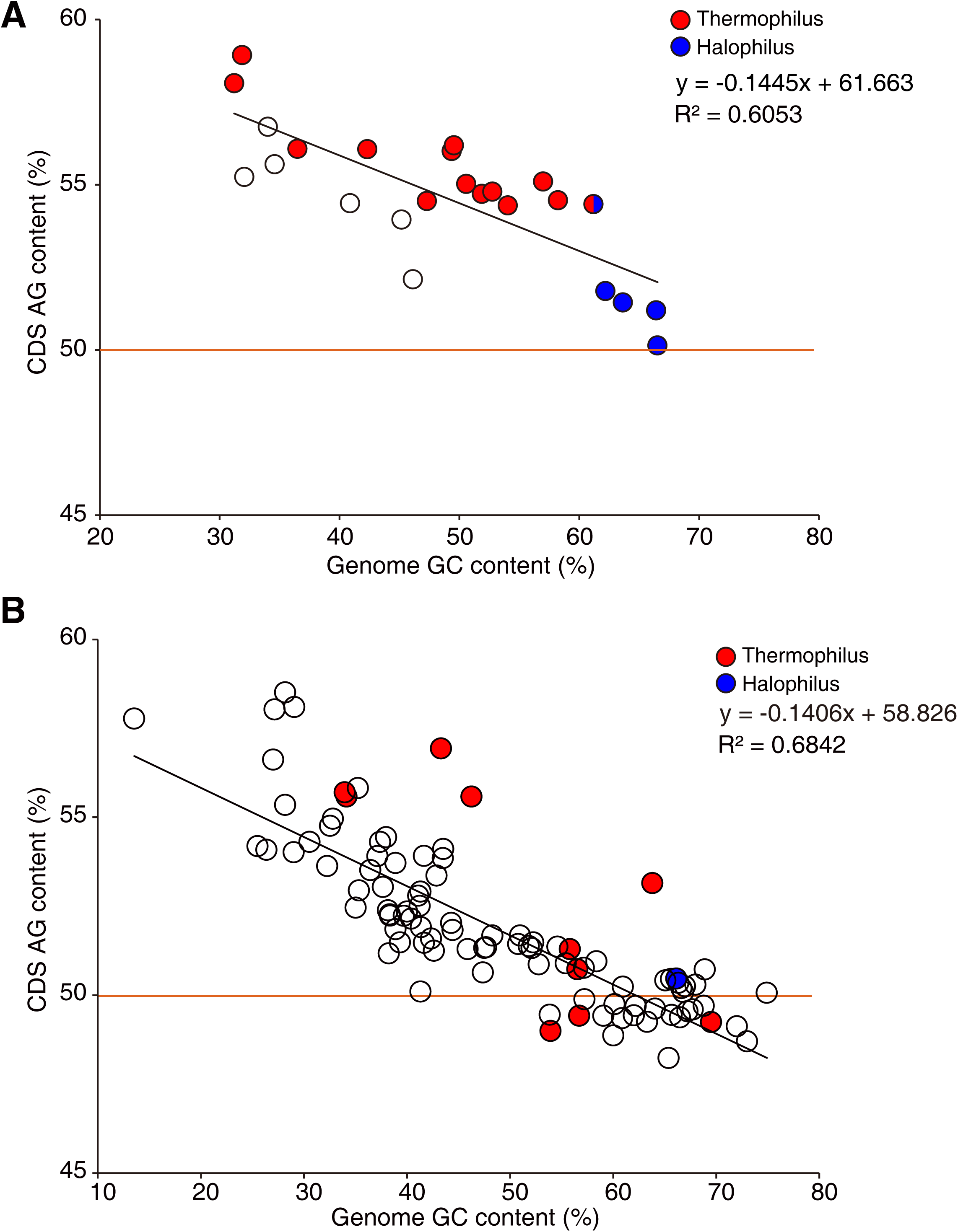
Correlation between CDS purine contents and genome GC contents in archaea (A) and bacteria (B). Each point is one species.

**Fig. S5.**
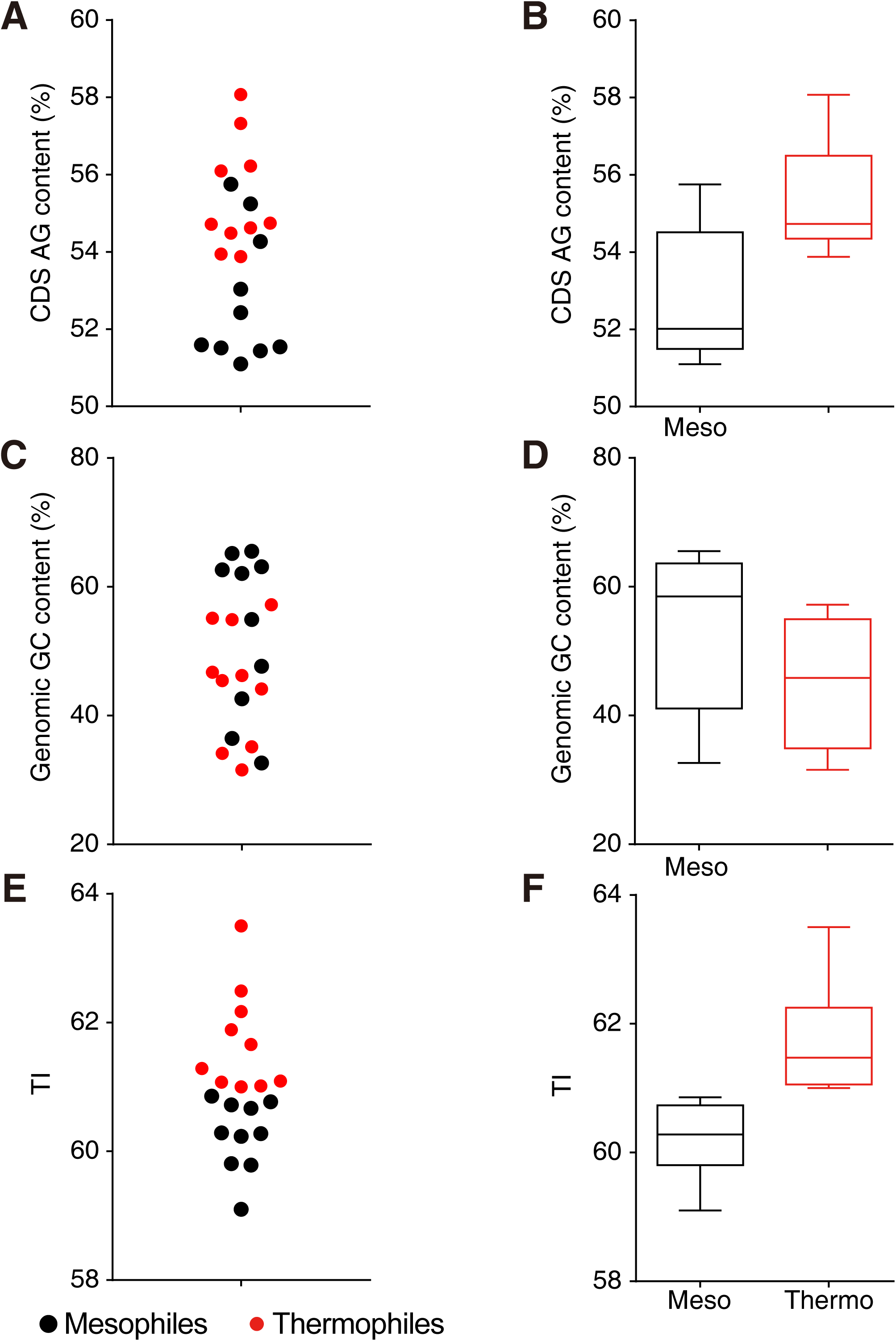
CDS purine contents, genome GC contents and *TI* of archaea. 10 thermophilic and 10 mesophilic archaea were analyzed (Table S3). The CDS purine contents, genome GC contents and *TI* were plotted as scattered plots (A, C, E) and box and whiskers charts (B, D, F).

**Fig. S6.**
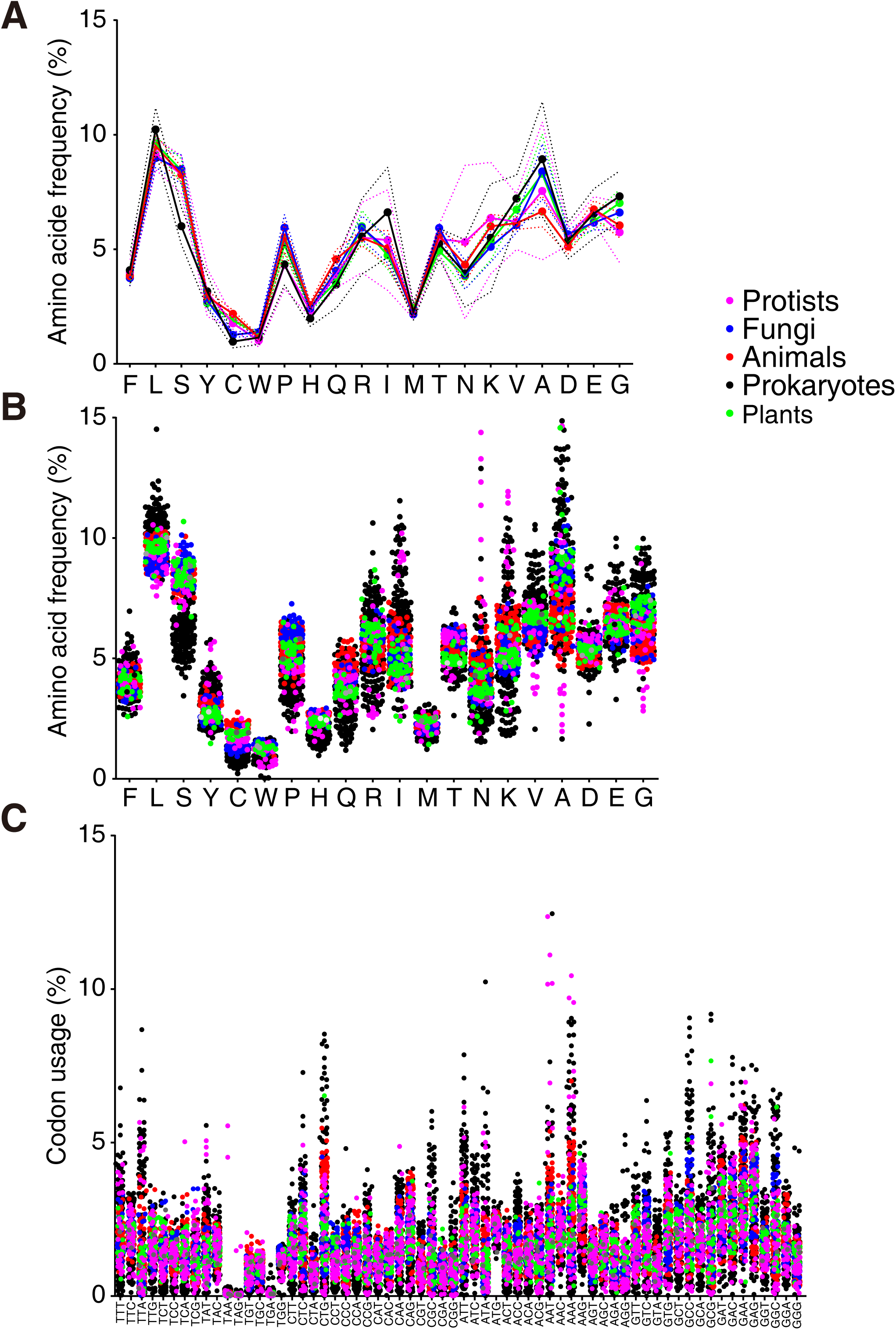
Amino acid frequencies and codon usages of all 371 genomes. A, average amino acid usages were taken by groups, dotted lines representing standard deviations; B, C, amino acid frequencies (B) and codon usages (C) of all 371 genomes, each point representing one genome.

**Fig. S7.**
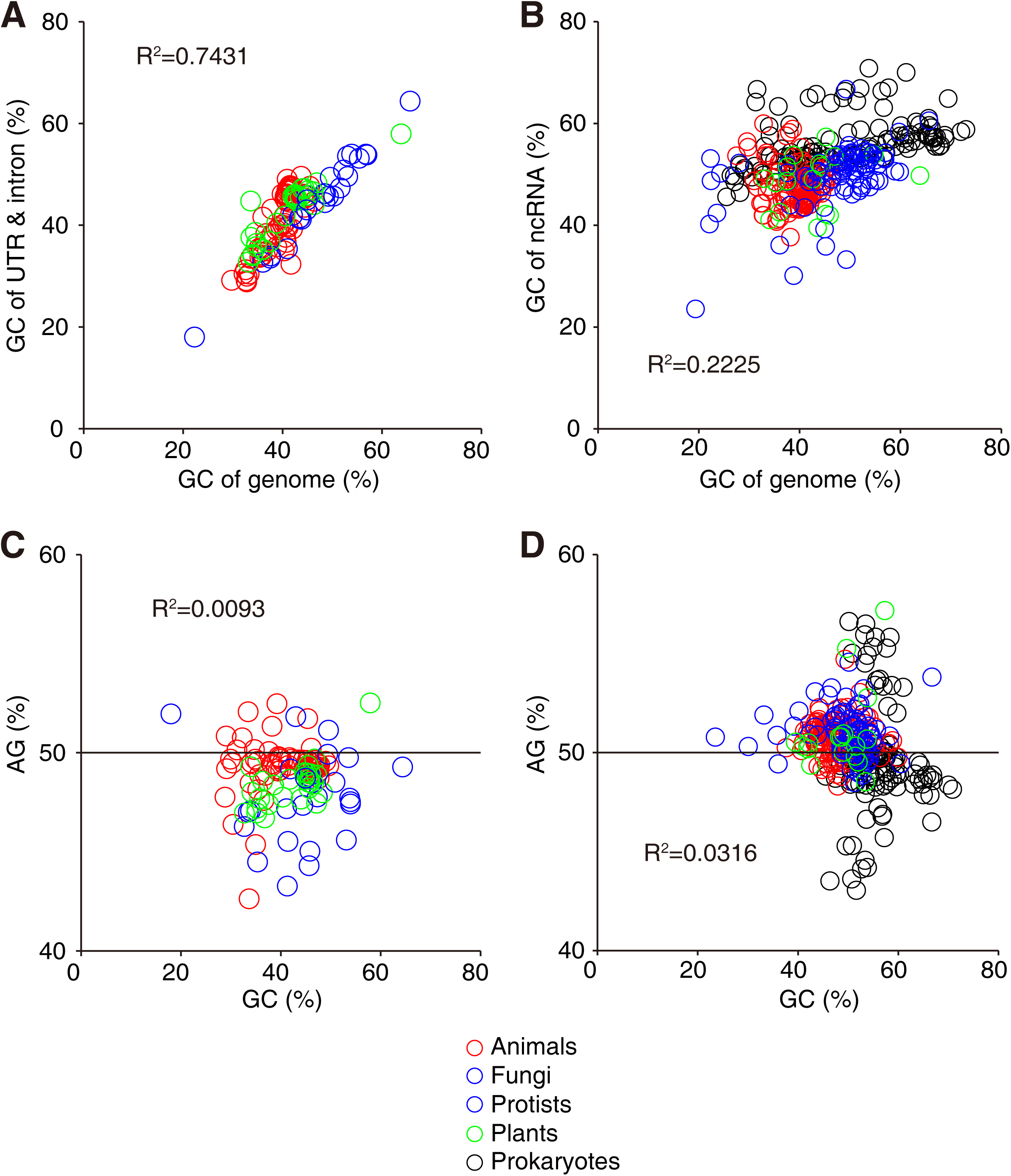
Nucleotide compositions of UTR, introns and noncoding RNAs. A, B, the correlations of GC contents between UTR/introns and genomes (A), and between noncoding RNA genes and genomes (B); C, D, AG-GC plots of UTR/introns (C) and noncoding RNA genes (D).

**Fig. S8.**
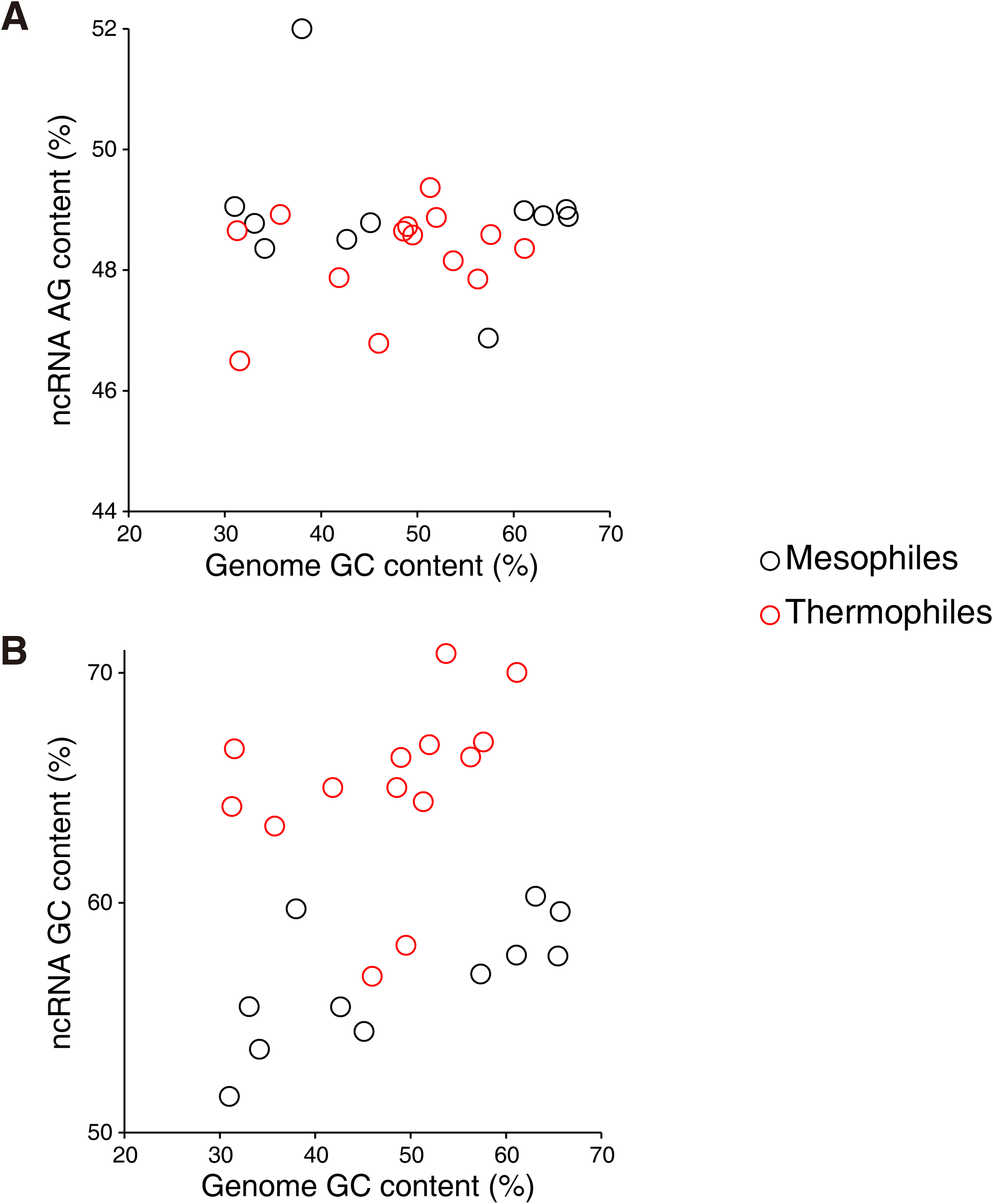
Purine contents and GC contents of ncRNAs in archaea. No correlations were found between ncRNA purine content and genome GC content (A), or between ncRNA GC content and genome GC content (B). The ncRNA GC contents of thermophiles are significantly higher than those of mesophiles (B).

